# Molecular annotation of AOPs guides the development of the next generation mechanistic chemical safety assessment and new approach methods

**DOI:** 10.1101/2022.07.08.499301

**Authors:** Laura Aliisa Saarimäki, Jack Morikka, Alisa Pavel, Seela Korpilähde, Giusy del Giudice, Antonio Federico, Michele Fratello, Angela Serra, Dario Greco

## Abstract

Mechanistic toxicology has emerged as a powerful framework to inform on the safety of chemicals and guide the development of new safe-by-design compounds. Although toxicogenomics provides support towards mechanistic evaluation of chemical exposures, the implementation of toxicogenomics-based evidence in the regulatory setting is still hindered by uncertainties related to the analysis and interpretation of such data. Adverse Outcome Pathways (AOPs) are multi-scale models that link chemical exposures to adverse outcomes through causal cascades of key events (KEs). The use of mechanistic evidence through the AOP framework is actively promoted for the development of new approach methods (NAMs) and to reduce animal experimentation. However, in order to unleash the full potential of AOPs and build confidence into toxicogenomics, robust and unified associations between KEs and patterns of molecular alteration need to be established.

Here, we hypothesised that systematic curation of molecular events associated with KEs would enable the modelling of AOPs through gene-level data, creating the much-needed link between toxicogenomics and the systemic mechanisms depicted by the AOPs. This, in turn, introduces novel ways of benefitting from the AOP concept, including predictive models, read-across, and targeted assays, while also reducing the need for multiple testing strategies. Hence, we developed a multi-step strategy to annotate the AOPs relevant to human health risk assessment. We show that our framework successfully highlights relevant adverse outcomes for chemical exposures with strong *in vitro* and *in vivo* convergence, supporting chemical grouping and other data-driven approaches. Finally, we defined and experimentally validated a panel of robust AOP-derived *in vitro* biomarkers for pulmonary fibrosis.

## Introduction

Mechanistic aspects of chemical exposures have been long exploited in the context of academic research, resulting in the emergence of toxicogenomics and systems toxicology as independent fields (1,2). Although the mechanistic insight gained through the technologies employed in academia has been valued as supporting evidence in the regulatory setting, its incorporation into the regulatory framework is to date hindered by concerns related to the robustness and reproducibility of such data and its analysis (3). At the same time, the growing need for faster, cheaper, and more ethical approaches for chemical safety assessment have made mechanistic toxicology central for clarifying aspects important to regulatory decision making. Furthermore, uncovering exposure related mechanistic properties is emerging as a fundamental approach for the design of new drugs and chemicals (4,5). Hence, multiple high-end research initiatives are underway to drive the shift from traditional animal-based assessment of apical toxicity endpoints towards *in vitro* and *in silico* approaches supported by mechanistic evidence (6-8).

Adverse Outcome Pathways (AOP) emerged as models to organise biological mechanisms into causally linked sequences of multi-scale events to support chemical risk assessment (9). AOPs have since expanded beyond the limits of toxicology, showing their applicability in organising mechanisms of disease progression and adverse health outcomes (10,11), and could even be applied to assess beneficial effects of therapies. The mechanisms depicted by AOPs comprise a sequence of events that progress from the molecular initiating event (MIE) towards an adverse outcome (AO) through intermediate steps, key events (KEs), with biological complexity increasing as the AOP progresses. Individual KEs are connected by key event relationships (KER), that verbally explain the causal link between the events and provide context for the pathway.

The AOP concept quickly attracted attention due to its potential in tackling one of the major challenges in the shift away from traditional toxicology: deciphering systemic and long-term outcomes of chemical exposures without the use of animal experiments. While significant efforts still need to be made towards this goal, AOPs encompass the means to systematically guide the integration of *in vitro* -based evidence into the risk assessment framework (12). AOPs provide the grounds for various predictive approaches, read-across, and the development of targeted assays and new approach methods (NAMs), as also suggested by regulatory agencies and international organisations, such as the OECD (8). Furthermore, the construction of AOPs can help identify gaps in knowledge and guide resources towards mechanisms in need of further investigation, or alternatively, reveal connections that have not been previously characterised (13).

Concurrently with the development of the AOP framework, the role of omics data in elucidating biomarkers and mechanisms of action of chemical exposures and diseases has become more prominent (14-18). Omics data have been used to support the development of AOPs, especially through the identification of molecular targets and mechanisms (19-23). However, full exploitation of omics-based evidence in the context of AOPs is hindered by the complication of linking molecular data to complex biological events, affecting both the development and the application of AOPs. Furthermore, while the value of omics data in answering questions of regulatory importance is recognised, the lack of standardisation in analysis and reporting have hampered widespread regulatory acceptance of omics-based evidence (24). Bypassing these challenges could broaden the application of AOPs and further aid in the development of the concept towards quantitative models and assays. While molecular assays based on arbitrarily selected reporter genes have been proposed (e.g., ToxCast assays), there is an urgent need to develop new data-driven unbiased molecular assays for reliable and efficient mechanistic safety assessment of chemicals. Here we hypothesised that rigorous curation of molecular events associated with AOPs could facilitate the implementation of omics-based evidence to 1) support the development of new AOPs, 2) identify and fill gaps in knowledge, and 3) transfer AOP-based knowledge into robust assays to support chemical safety assessment.

Well-curated gene ontologies, pathways, and biological processes are used to interpret omics results and their translation into biologically relevant information. While some KEs can be easily crosslinked with such terms and their associated genes, the annotation of complex KEs taking place at a higher level of biological organisation (e.g., at the tissue- or organism-level) is a more demanding task. This requires knowledge regarding ontologies and the biological events themselves. For instance, generic annotations are helpful for categorising KEs, but without the intention of modelling KEs using the associated gene sets, they will likely not reach the level of granularity required for such a task. This is currently reflected in the annotations provided in the AOP-Wiki repository (aopwiki.org). The annotation of KEs to selected ontologies is currently included as an option in AOP-Wiki. However, the coverage of the annotations is currently low and has not been intended for modelling the KEs using the gene sets associated with their annotations.

Previous efforts to curate external annotation have shown the potential of the approach (25,26). However, these have either remained at the level of abstract associations or focused on individual examples (27,28). Hence, systematic, fit-for-purpose, and up-to-date annotation linking KEs to curated gene sets has not yet been established. To this end, we applied an integrated strategy for defining gene-KE-AOP associations through systematic curation. We show the applicability of our strategy for evaluating potential adverse outcomes of chemical exposures, and for the identification of AOP-driven biomarkers that can inform the development of target assays and novel approaches to chemical hazard characterisation.

## Results and discussion

We developed an integrated approach to systematically associate curated gene sets to the KEs and AOPs. Our approach combines natural language processing (NLP) techniques with manual curation to link relevant biological processes and pathways, as well as their associated genes, to KEs of AOPs relevant for human health risk assessment. The resulting gene-KE-AOP connections enable the modelling of KEs and AOPs through gene-level data, which further introduces novel ways to benefit from the AOP concept. We applied this approach to generate an AOP fingerprint for a known profibrotic exposure *in vivo* and *in vitro* and finally combined the annotation to a framework for prioritising KE- and AOP-associated genes to guide the discovery of biomarkers and reporter genes. The complete approach described in the following sections is summarised in Figure 1.

**Figure 1.**
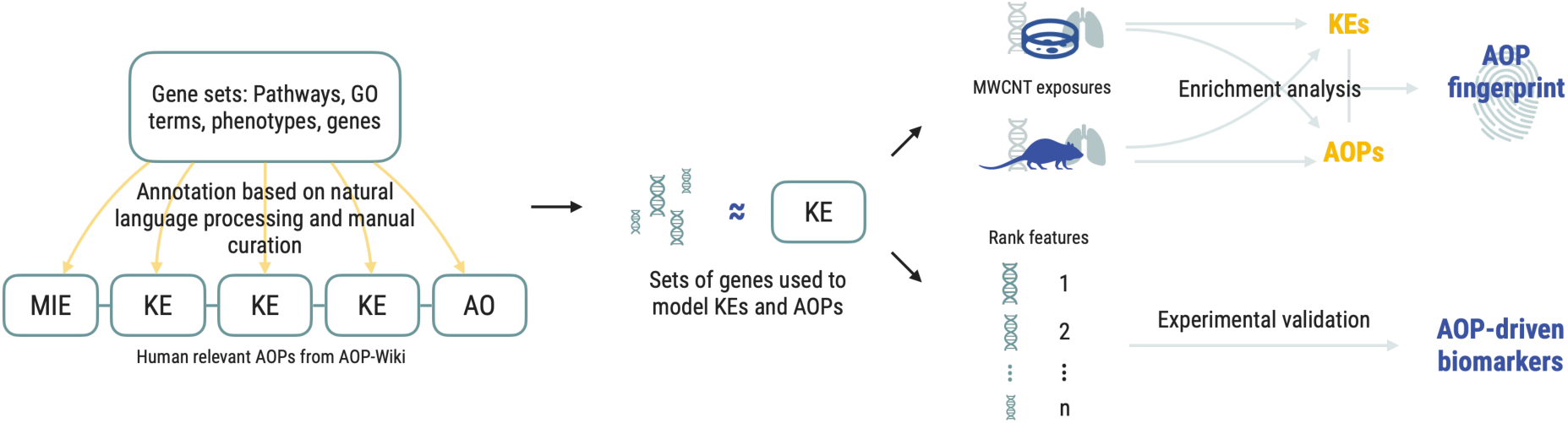
Overall scheme of the study. Established gene sets were annotated to KEs of the AOPs relevant for human health risk assessment. The resulting gene sets were then used to model the KEs. The validity of the annotation was evaluated using gene signatures of exposures with known adverse outcomes. Finally, we combined the approach with a gene prioritisation framework resulting in the identification of AOP-driven biomarkers for pulmonary fibrosis.

### The majority of KEs can be successfully annotated to curated gene sets

At the time of retrieving the data from the AOP-Wiki repository (November 2020), a total of 289 AOPs and 1,131 distinct KEs were identified. However, after eliminating the AOPs for which taxonomic applicability was either not available nor in the scope of our investigation, 176 AOPs and 856 unique KEs remained, forming a total of 1,245 unique AOP-KE pairs (specific KEs). Although the AOP-Wiki houses selected annotations for some of the KEs, majority of them were considered not to be specific enough for our purpose (i.e., KEs describing the dysregulation of a specific gene annotated to terms such as “gene expression”). Additionally, as the existing annotations only cover a part of the KEs, we decided to consistently curate the annotation of all KEs. As a result, 799 unique KEs mapped to 175 AOPs received a curated annotation. The KEs were treated as individual instances, hence the same KE mapped to multiple AOPs was always annotated to the same term(s). A summary of the number of terms annotated to the KEs is presented in Figure 2A along with the proportions of the different term sources (Figure 2B). GO biological processes (GO_BP) represent most of the mapped annotations, followed by GO molecular functions (GO_MF) and Human Phenotype Ontology (HPO). Since up to five annotations were provided for the KEs, the final gene sets used from herein comprise the union of the genes mapped to each annotated term. This structure allowed improved specificity, while also providing the possibility to further refine the gene sets using the hierarchical order implemented where applicable. The size of the gene sets associated to each KE range from one to 5,990 genes, with a median value of 81 genes. Consequently, when AOPs are modelled by combining the gene sets associated to their KEs, the gene set sizes range from 15 to 5,992 with the median size being 752 genes.

**Figure 2:**
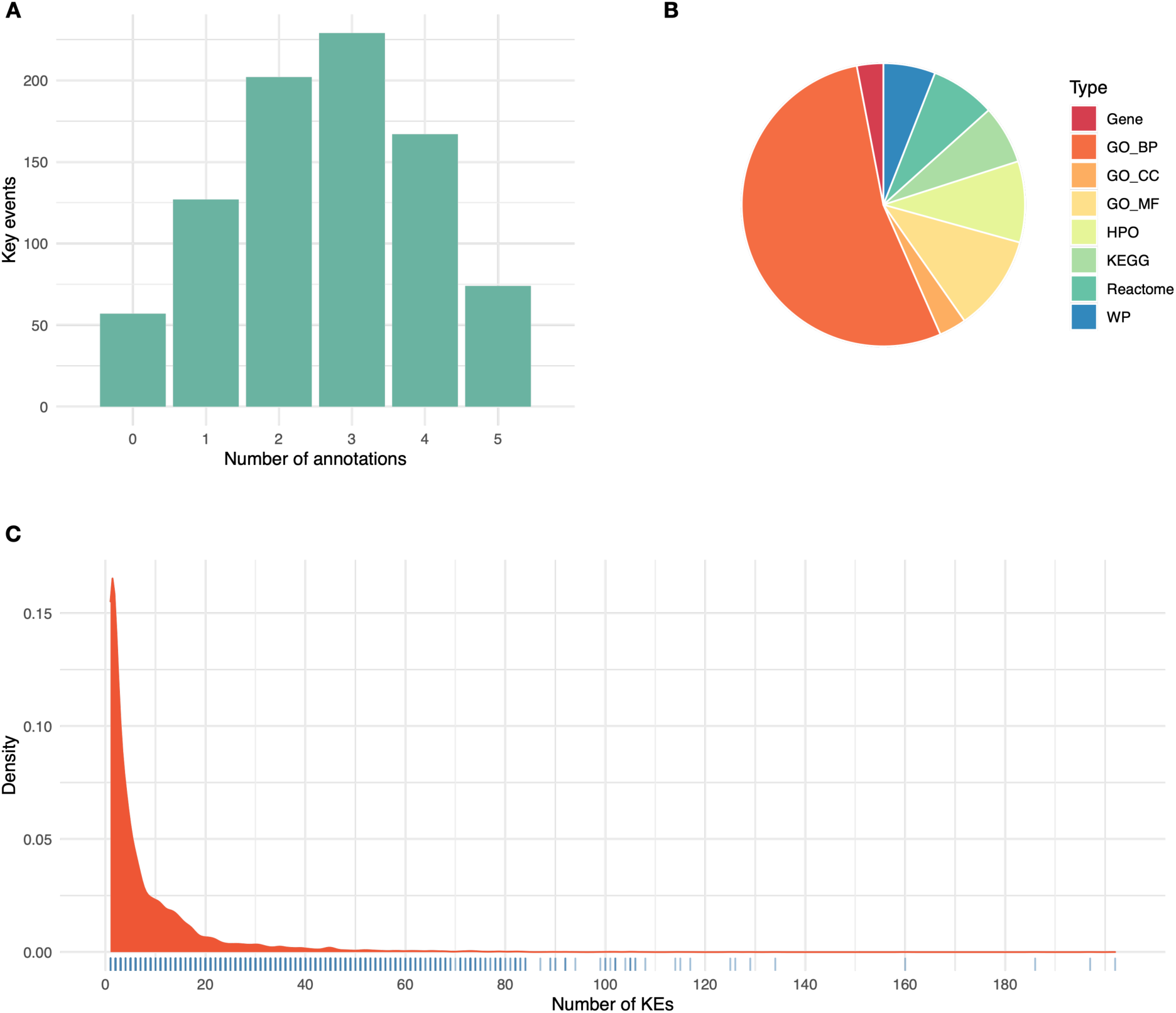
Descriptive analysis of the KE annotation. (A) Bar plot describing the number of annotated terms per KEs. (B) Pie chart expressing the proportions of different annotation types. (C) Density distribution of the number of KEs each gene is annotated to.

In total, the annotations comprise 15,825 genes. While the majority of genes are annotated to less than 5 KEs (9,044 genes), 1,434 genes have more than 20 KEs associated to them, and 50 genes have more than 80 associated KEs (Figure 2C). Although these numbers can be affected by annotation bias, e.g., certain genes are better researched and annotated than others, they can also guide the selection of AOP-driven biomarkers when specificity is of importance.

The result of the annotation suggests that the majority of human relevant KEs can be linked to gene ontology, phenotype, and pathway terms, and further to their corresponding sets of genes. Previous efforts to annotate KEs have provided similar implications (25,26). However, the applicability of the annotations has not been previously systematically explored, nor have they allowed modelling of the KEs and AOPs by use of gene sets on a large scale. Here, we present the first thorough mapping of all human relevant KEs to curated gene sets.

### AOP enrichment highlights relevant adverse outcomes associated to chemicals

We tested the ability of our novel annotations to highlight relevant AOPs by analysing a set of curated reference chemicals as defined by EU Reference Laboratory for alternatives to animal testing (ECVAM) and National Toxicology Program Interagency Center for the Evaluation of Alternative Toxicological Methods (NICEATM). We focused on four categories of chemicals defined by their toxicity properties to include hepatotoxic and carcinogenic agents as well as thyroid disruptors and sex hormone receptor (estrogen receptor – ER, and androgen receptor - AR) agonists. For each of the selected chemicals, we retrieved a list of associated genes from the Comparative Toxicogenomics Database (CTD) (29), resulting in a final set of 75 chemicals (Supplementary File 1).

AOPs related to each category were then identified among the 175 AOPs we had curated, and the prevalence of relevant AOPs (i.e., AOPs describing carcinogenesis for carcinogenic chemicals, etc.) among the five most significantly enriched AOPs for each chemical were evaluated. The results suggest that the enrichment approach successfully highlights AOPs of relevance for each group of chemicals (Figure 3). All sex hormone receptor agonists had at least one relevant AOP among the top five enriched, while the proportions varied from 43% (thyroid disrupters) to 93% (carcinogens) in the other categories (Figure 3).

**Figure 3:**
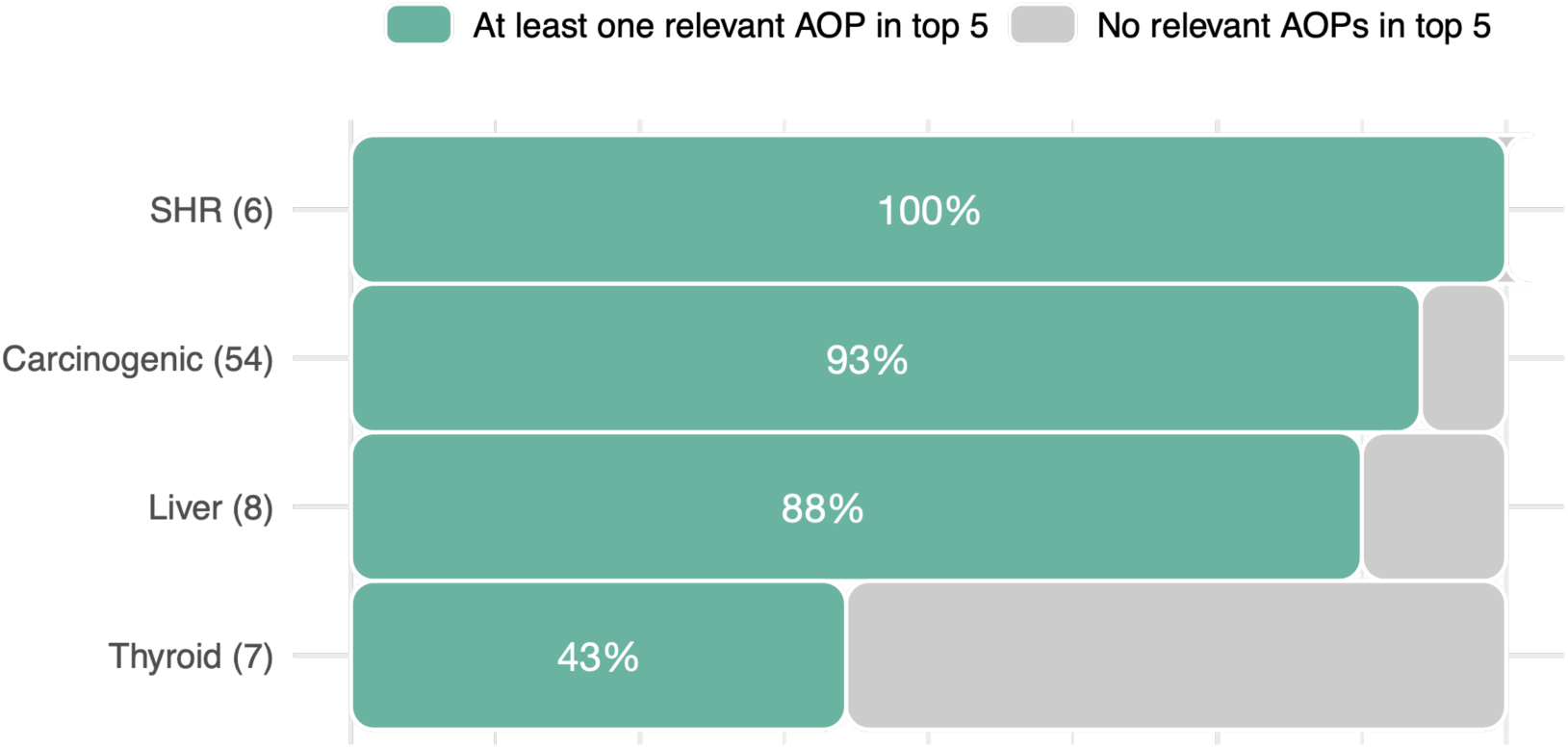
Bar plot representing the proportion of chemicals with relevant AOPs among the top five enriched AOPs based on the chemical classification. Number in brackets after the category name refers to the number chemicals in each category while the percentage on the bars reflects the proportion of chemicals in each category highlighting relevant AOPs. SHR stands for sex hormone receptor agonist.

In the group of carcinogenic chemicals, 93% of the compounds evaluated had cancer-related adverse outcomes among the top enriched AOPs. In fact, the group of carcinogens had the highest proportion of relevant AOPs at the top as compared to the others (median four out of five compared to the median of two out of five in the other groups). However, it should be noted that AOPs related to cancer are among the most represented group of AOPs, and cancer-related genes are generally highly researched and annotated, which may introduce a level of annotation bias that should be recognised.

The remaining four carcinogenic chemicals (7%) that showed no cancer AOPs among the top enriched AOPs were N-nitrosodiethanolamine, N-nitrosomorpholine, phenacetin, and tetrachloroethylene. N-nitrosomorpholine and N-nitrosodiethanolamine are both nitrosamines whose suspected adverse outcomes besides carcinogenesis include non-alcoholic steatohepatitis (30). Indeed, both compounds contained hepatic steatosis related AOPs among the top five enriched AOPs (Supplementary File 1). Tetrachloroethylene (perchloroethylene, PCE) is a chlorocarbon solvent used in dry-cleaning and other degreasing applications (31). AOPs with the most significant enrichment for PCE were also related to hepatic adverse outcomes. Although neurotoxicity is one of the most frequent AOs associated with PCE exposure, hepatotoxicity has also been reported (31). Our results documenting liver steatosis are supported by biopsy-based evidence of liver disease, both in human and animal models, in settings of high occupational exposures (32). Lastly, phenacetin is a drug that was widely used as pain medication until it was withdrawn from the market across the globe due to increasing evidence of carcinogenicity and renal toxicity (33). The most enriched AOPs for phenacetin included immune related AOPs “Immune mediated hepatitis” (Aop:362) and Aop:277 titled “Inhibition of IL-1 binding to IL-1 receptor leading to increases susceptibility to infection”. Although there is no described association between phenacetin and IL-1 or immuno-toxicity, it is known that they both play a role in paracetamol-associated liver toxicity, which is the main metabolite of phenacetin (33,34).

In the case of the known liver toxicants, hexaconazole was the only compound not highlighting AOPs associated with liver toxicity among the top enriched AOPs. Hexaconazole is a widely used triazole fungicide. It acts by blocking sterol biosynthesis *via* inhibition of cytochrome P450 (35). Hexaconazole was considered as a Group C-Possible Human Carcinogen by the US EPA due to increased incidence of benign Leydig cell tumours in rats (https://www3.epa.gov/pesticides/chem_search/hhbp/R000356.pdf). Moreover, it was found to affect the reproduction of female rats (35). The top enriched AOPs correctly identify this signature. Furthermore, the top two pathways “HMG-CoA reductase inhibition leading to decreased fertility” and “Modulation of Adult Leydig Cell Function Subsequent to Decreased Cholesterol Synthesis or Transport in the Adult Leydig Cell” both suggest a decrease in cholesterol levels by inhibition of the HMG-CoA reductase. Drugs inhibiting this enzyme, such as statins, are known to possibly cause liver damage (36).

Known thyroid toxicants performed poorest in our analysis. Bifenthrin, malathion, permethrin and simazine did not capture thyroid-related AOPs among the top five enriched. All these compounds have been widely used in agriculture as herbicides or pesticides. Agrochemicals represent a significant class of endocrine disrupting chemicals, albeit through varying mechanims. It is now accepted that many of these molecules may mimic the interaction of endogenous hormones with nuclear receptors, such as estrogen, androgen, and thyroid hormone receptors (37). Indeed, bifenthrin has already been reported as an endocrine-disrupting compound by blocking the binding of endogenous hormones (38). In our framework, its anti-estrogenic activity emerges as the most enriched AOP (Supplementary File 1). Malathion is an organophosphate pesticide that is known for its low acute toxicity and rapid degradation (39). In this light, it is not listed as a primary thyroid disrupting chemical, and its toxicity has been associated with the inhibition of acetylcholinesterase activity on nerve impulse (39). Recent studies, however, demonstrated that malathion acts as an endocrine disruptor, both *in vitro* and *in vivo* (40,41). Our results support these findings, highlighting the Aop:165: “Anti-estrogen activity leading to ovarian adenomas and granular cell tumors in the mouse” as well as Aop:112: “Increased dopaminergic activity leading to endometrial adenocarcinomas“. Furthermore, Moore *et al*. demonstrated that malathion exposure at higher concentrations induces cytotoxic and genotoxic effects in HepG2 through oxidative stress, which can finally lead to liver cancer (39). Similarly, our framework highlights both the “PPARalpha-dependent liver cancer” and “Cyp2E1 Activation Leading to Liver Cancer” AOPs. Simazine is a triazine herbicide whose use has been banned in most European countries for nearly two decades (42). Simazine has now been recognised, similarly to the other compounds, as an endocrine disrupter (42). Interestingly, the enrichment analysis for simazine highlighted AOPs related to the development of adenomas and carcinomas through endocrine disrupting activities (e.g., Aop:107 titled “Constitutive androstane receptor activation leading to hepatocellular adenomas and carcinomas in the mouse and the rat”) as well as direct disruption of the GnRH pulse (Supplementary File 1). Although multiple *in vivo* and *in silico* evidence also indicate permethrin as possible endocrine disruptor (43,44), no endocrine related pathways are present in the top enriched AOPs. However, this framework was able to highlight the modulating effect of permethrin on the lipid metabolism. It has been demonstrated that in HepG2 cells, permethrin increases lipogenesis and decreases beta oxidation, possibly contributing to the development of NAFLD (45). Indeed, the “Inhibition of fatty acid beta oxidation leading to nonalcoholic steatohepatitis (NASH)” AOP is statistically enriched in our results.

Together, these results highlight relevant AOPs modelled by our curated gene sets to be enriched by the genes associated to the compounds, suggesting that our framework is able to support robust mechanistic and data-driven chemical grouping as well as the identification of potential adverse outcomes using chemical-gene associations.

### Our annotation enables grouping of KEs resulting in improved modelling of the AOP network

In order to fully unleash the potential of mechanistic toxicology, more informative testing strategies need to be developed that can monitor specific phases of the exposure-bio interactions and mechanisms. To this end, we defined accurate sets of genes capable of modelling specific KEs and AOPs. However, one of the challenges observed in the AOP-Wiki is the redundant semantics in the naming of KEs. While creating a new KE can be meaningful in many cases (e.g., the same biological process taking place in a distinct organ or a tissue), unnecessary redundancy can lead to challenges in the application of the AOP-based knowledge. This is especially true when modelling AOPs as a network and using such representation to identify hidden connections and to perform read across analysis (10,46–51).

Hence, we hypothesised that KEs could be grouped based on the degree of similarity of their associated gene lists. We calculated the similarity of the KEs based on their annotated gene sets and grouped together those mapped to identical sets of genes (Jaccard Index = 1). This resulted in the identification of 637 groups of varying sizes. These groups were characterised by four main concepts: 1) truly duplicated KEs due to distinct semantics, 2) same biological event in multiple biological systems, 3) subsequent KEs mapped to the same terms due to inadequate specificity, and 4) opposite regulation of the same biological event (i.e., increased *vs*. decreased signalling).

Here, the grouping based on identical gene sets was selected due to the nature of the downstream application and statistical considerations (i.e., to avoid multiple testing against the same gene set in enrichment analysis). However, a parallel approach with varying cut-off values for similarity could be implemented to cluster KEs more roughly and to define specific categories of events. Similarly, further refinement of the KE clusters could help to enhance the AOP network by removing redundant nodes which, in turn, could reveal hidden links.

The potential of the KE grouping was showcased using a subgraph formed by considering the AOPs related to pulmonary fibrosis (PF). PF is a chronic lung disease characterised by tissue damage and scarring that impairs lung function (52). A range of environmental exposures, including certain chemicals, drugs, nanomaterials and radiation, as well as infectious diseases have been identified as causative agents for PF (52-54). Moreover, the COVID-19 pandemic has raised concerns about increasing rates of PF (55-57). Understanding the disease mechanisms can help in the development of strategies to treat and prevent the disease, and to control and modulate the exposures that contribute to its progression. Furthermore, it can serve as the foundation for developing targeted assays for evaluating PF as a toxicological endpoint.

Six AOPs related to PF were available in the AOP-Wiki at the time of data retrieval (Figure 4A). These distinct AOPs characterise multiple pathways leading to the same adverse outcome. Together, these AOPs comprise 30 KEs, which form a connected graph when modelled as a network (Figure 4C). However, several redundancies were observed among the KEs. For instance, the adverse outcome was expressed either as *lung fibrosis* (Event:1276) or *pulmonary fibrosis* (Event:1458). Hence, the application of the similarity-based grouping resulted in 23 distinct KEs (Figure 4B) that were then used as the basis for merging the KE nodes in the PF network (Figure 4D).

**Figure 4:**
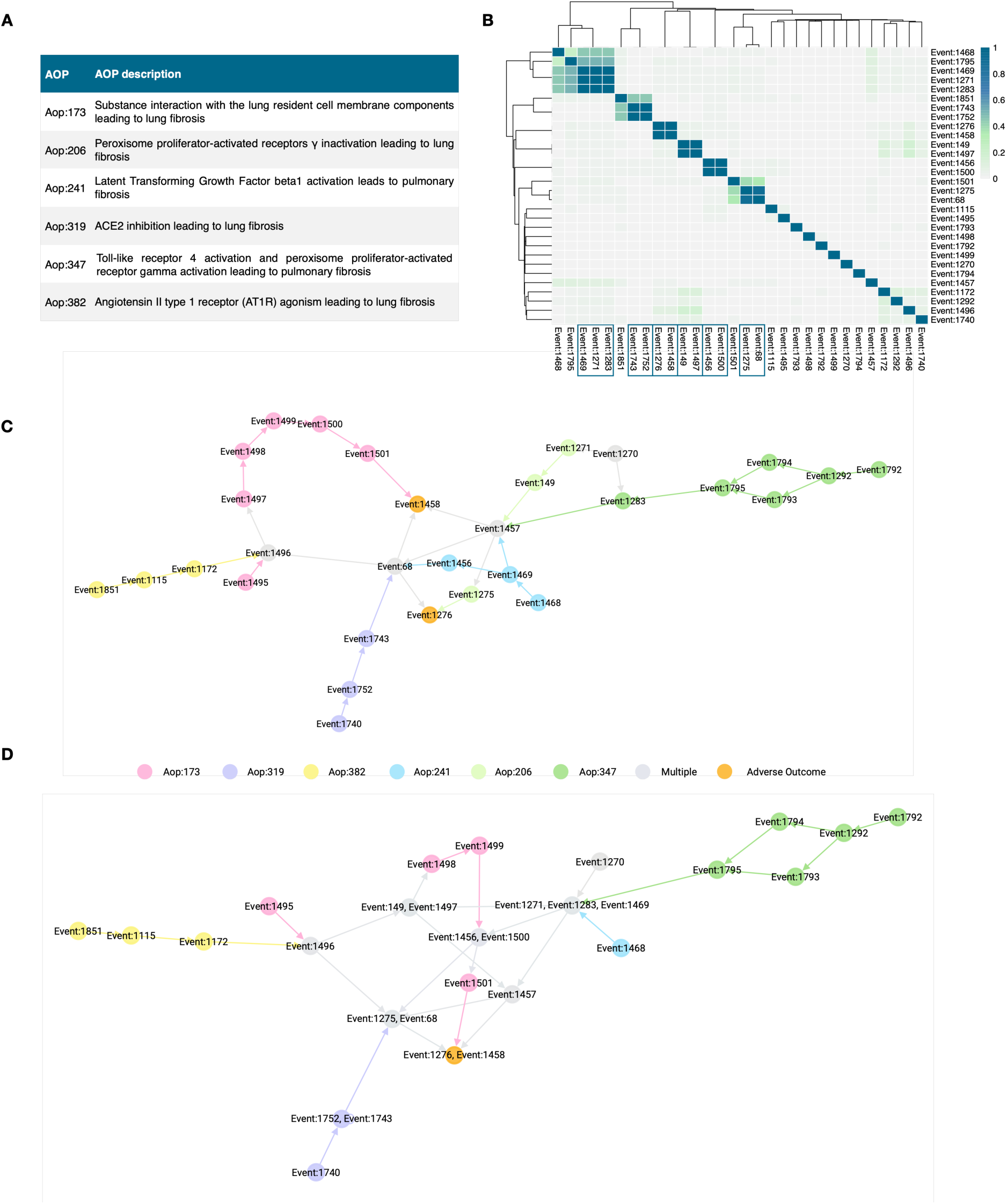
A) Table presentation of pulmonary fibrosis (PF) AOPs identified in the current study. B) Heatmap representing the Jaccard index-based similarity of the PF KEs as per their associated gene sets. Values close to zero (light grey) correspond to a low similarity between distinct KEs, while the increasing similarity is expressed with the colour changing through green to blue. C) Graph presentation of the PF AOPs using their original KEs. Distinct colours denote the KEs of individual AOPs, grey nodes are KEs shared by multiple AOPs, and orange nodes correspond to the shared adverse outcomes. D) Graph presentation of the PF AOPs after KE grouping. The number of shared (grey) nodes has now increased, and the duplicated AO has been grouped into one distinct AO (orange).

The PF AOPs formed a connected network, indicating that each of the individual AOPs shared at least one KE with one or more of the other AOPs. However, as the duplicated KEs were merged, the similarities between the AOPs became more evident. This is evidenced by the increasing number of shared KEs in the graph after merging (the grey nodes in Figure 4D) as compared to the original graph (Figure 4C). Furthermore, the merging revealed Aop:206 to be fully contained within the other AOPs.

The refinement of the AOP network through KE grouping simplifies the network while also enhancing the robustness of the KE relationships, depicted by the connections between the nodes. This process, in fact, removes redundant nodes, which supports the application of AOP networks in AOP research and risk assessment. Furthermore, as duplicated events are removed, the true influence of each node can be assessed more robustly though network analytics.

This example demonstrates the effect of KE redundancy and the potential of data-driven grouping of the KEs. While manual assessment and grouping would be achievable for a limited number of AOPs at a time, doing it AOP-Wiki wide would be a massive undertaking. Here, we show how our curated gene-KE-AOP connections can help guide the grouping and hence enhance network-based approaches in AOP research. Furthermore, our results suggest that it is often possible to identify meaningful genes that can successfully monitor multiple key events of a similar nature.

### The AOP fingerprint of multi-walled carbon nanotubes converges *in vitro* and *in vivo*

Toxicogenomics has supported the development of mechanistic toxicology and further enhanced the possibility to obtain relevant information from *in vitro* studies, which could reduce the need for animal experimentation (58-60). Here, we tested the hypothesis that toxicogenomic data generated in two independent *in vitro* and *in vivo* exposure models would converge on a robust set of relevant AOPs. We focused on Mitsui-7, a known hazardous long and rigid multi-walled carbon nanotube (MWCNT). The airways provide the most prominent route of exposure to this nanomaterial, and it is best characterised for its lung-related adverse outcomes, including fibrosis (61-64). Hence, we selected data derived from a lung exposure to the MWCNT in mice (65), and an *in vitro* dataset with exposures on four cell lines representative of the human airways (59,66). These cell lines include differentiated THP-1 cells as a model of macrophages, A549 representing alveolar basal epithelial cells, BEAS-2B as bronchial epithelial cells, and MRC as a model of lung fibroblasts. Differentially expressed genes (DEGs) from all experimental conditions *in vivo* and each cell line *in vitro* were obtained from Saarimäki et al. (67) and merged into a single mechanism of action *in vivo* and *in vitro*, respectively.

We then performed enrichment analysis against both the AOPs and the KEs separately in order to evaluate the coverage of distinct KEs. We used the proportion of significantly enriched KEs to further filter the significant AOPs. This led us to identify 33 significant AOPs from the *in vivo* data, while 12 resulted significant from the *in vitro* exposure. These results were defined as the specific AOP fingerprint for the exposures, and it is presented in Figure 5.

**Figure 5:**
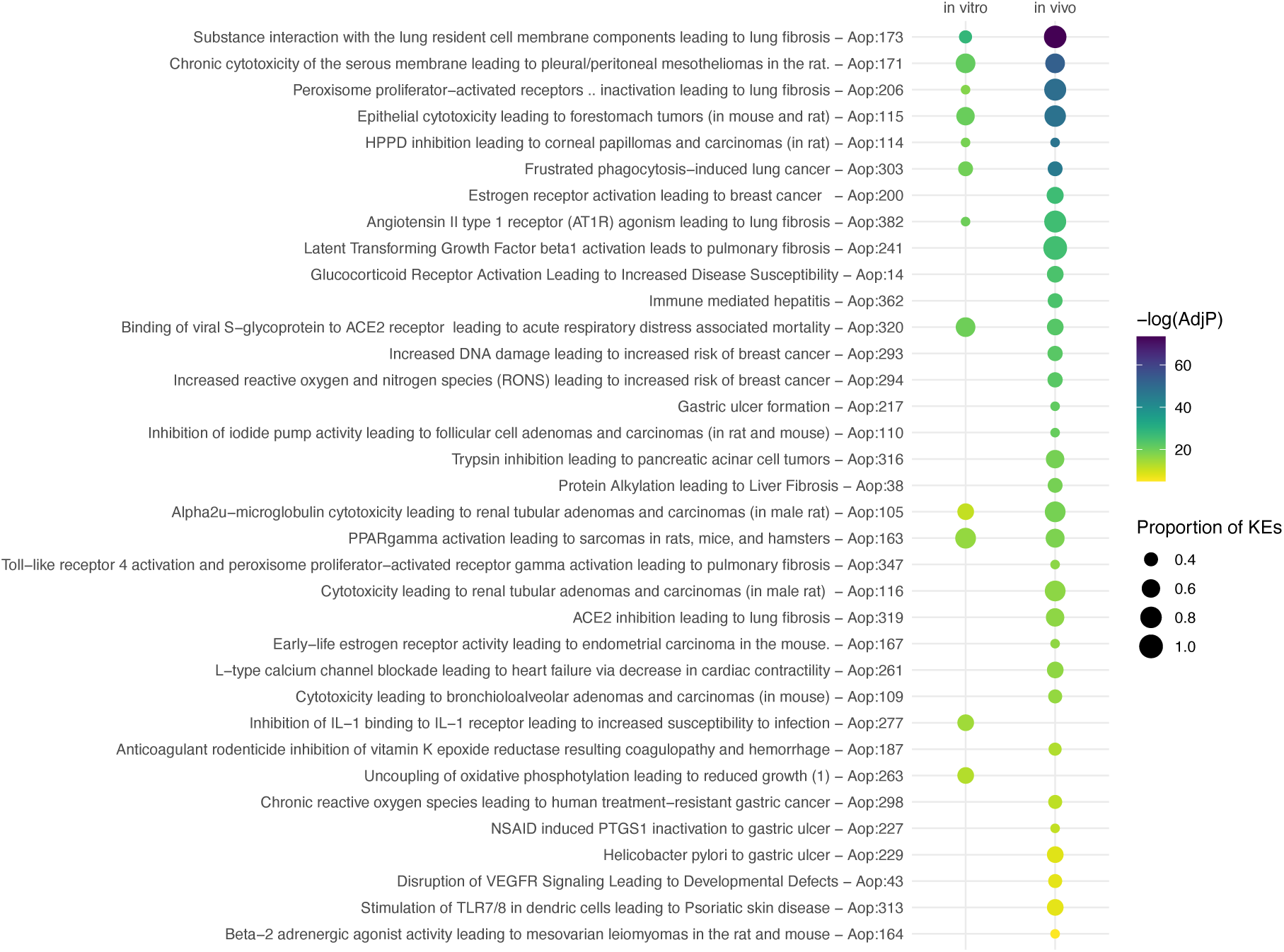
AOP fingerprint of Mitsui-7 exposure *in vitro* and *in vivo*. Size of the bubble reflects the proportion of significantly enriched KEs in an AOP, while to colour denotes the FDR-adjusted p-value in a negative logarithmic scale (i.e., the higher the number, the smaller the p-value). The AOPs are organised by the enrichment p-value from the *in vivo* data.

Despite the distinct sizes of the AOP fingerprints, 10 of the 12 AOPs enriched *in vitro* were also included in the *in vivo* fingerprint. Moreover, the top enriched AOPs were shared and ranked similarly between *in vivo* and *in vitro* when ranked by the smallest adjusted p-value. The AOP enriched with the most significant p-value in both instances was Aop:173 titled “Substance interaction with the lung resident cell membrane components leading to lung fibrosis” (Figure 5). The *in vivo* data set was able to capture 7 of the 8 KEs of the AOP as significantly enriched, while 3 out of the 8 KEs were enriched *in vitro*. Interestingly, Aop:173 has been specifically developed with robust evidence from MWCNT exposures, and multiple types of carbon nanotubes are listed as known stressors for the AOP (https://aopwiki.org/aops/173). The second AOP (Aop:171), on the other hand, describes the induction of pleural/peritoneal mesotheliomas by chronic cytotoxicity in rats. Like most AOPs used in this study, Aop:171 is still under development and lacks information on potential stressors. However, mesothelioma is a well-known adverse outcome of asbestos exposure, a fibrous silicate mineral whose adverse effects have often been used as a warning example for MWCNTs (68). Indeed, similarities in their mechanism of action have been extensively investigated (61,69,70).

The *in vitro* AOP fingerprint captures effects such as frustrated phagocytosis, oxidative stress, cytotoxicity, and immune activation, which have all been reported as consequences of this type of exposure and contribute to the pathogenic nature of Mitsui-7 (61,62,64,71).

Similarly, the profibrotic effects are highlighted with the multiple PF AOPs enriched. These effects are also observed in the *in vivo* AOP fingerprint. However, the *in vivo* fingerprint further highlights various AOPs outside the respiratory system, which is less apparent *in vitro*. While adverse outcomes beyond the immediate exposure site are feasible, many of these could likely be accounted for by the different effects of similar transcriptomic signatures in different biological systems (e.g., multiple AOPs related to gastric ulcer formation could reflect similar mechanisms of surfactant disturbance in two distinct exposure sites). On the other hand, the AOPs unique to the *in vitro* fingerprint, Aop:277 and Aop:263 (Figure 5), reflect the specific effects of the Mitsui-7 exposure on the immune system. Such specific signals can be easily masked in the *in vivo* system, where a large array of cell types is affected by the exposure.

It is worth noting that the exposures selected for the analysis had diverse set ups and a notable difference in the size of the combined mechanism of action (863 DEGs *in vitro vs.* 3,540 *in vivo*). While data from multiple cell lines were selected to capture effects besides immune cell activation *in vitro*, we were not able to match the dose and time course set up present in the *in vivo* dataset. However, we wanted to include this long-term exposure to evaluate whether it would result in broader coverage over the KEs of AOPs. Furthermore, histopathological evaluation from the same *in vivo* exposure set up has shown fibrosis in the lung from the day 7 onwards (72), suggesting that a whole PF AOP could be covered with this data. Indeed, all but the MIE (Event:1495) of Aop:173 were enriched *in vivo.* The high proportion of enriched KEs in the *in vivo* data supports the modelling of KEs with relevant gene sets and the use of toxicogenomic evidence for the development of AOPs, as well as the evaluation of potential adverse outcomes of chemical exposures. Likewise, we show that the analysis of toxicogenomic data against robustly annotated AOP framework supports a high degree of *in vitro* to *in vivo* extrapolation and further supports the inclusion of toxicogenomics-based evidence for regulatory purposes.

### KE-associated gene sets guide the selection of biomarkers

We showed that our KE-linked gene sets provide a robust way of evaluating potential outcomes of chemical exposures from transcriptomics data. This observation alone can help to guide chemical testing and grouping. However, to support the development of target assays and integrated approaches, specific reporter genes and markers need to be identified. Selection of transcriptional biomarkers and reporter genes only based on differential expression from experimental data gives little context or reference to the AO of interest. Even if a certain exposure is known to induce a specific endpoint, there is no indication whether the measured deregulation could be associated with the phenotype of interest. On the other hand, prioritising the features in the context of the KEs or whole AOPs could shed light on the importance and specificity of the feature regarding the phenotype. This, in turn, can guide the selection of potential biomarkers even in the absence of experimental data. Hence, we implemented a universal and customisable framework for the prioritisation of the KE-associated genes to drive the identification of AOP-informed biomarkers and used it to identify AOP-driven biomarkers for PF. The shortlisted marker genes were then screened by RT-qPCR in an *in vitro* model of human macrophages exposed to bleomycin, a well-known profibrotic chemical (73).

First, we defined characteristics for optimal biomarkers based on the Bradford Hill criteria, originally defined to evaluate causality in epidemiological research (74), but later adopted to other research fields as well (75). Our newly defined characteristics, their Bradford Hill counterparts, and short descriptions of the consideration of each step in the selection process are summarised in Table 1.

**Table 1:** We defined characteristics for optimal biomarkers based on the Bradford Hill criteria. The characteristics were then implemented into the prioritisation and selection protocol, and further to the evaluation of the prioritised genes.

| Bradford Hill | Our characteristic | Method/Assessment |
| --- | --- | --- |
| Consistency (reproducibility) | Reproducibility | Selection considers evidence from previous profibrotic exposures |
| Strength (effect size) | Amplitude | Significant alteration of the expression as compared to control |
| Experiment | Measurable | Transcriptional biomarkers measurable by qPCR; selected genes need to be expressed in the model |
| Biological gradient (dose-response relationship) | Dose-responsive | Benchmark-dose modelling to evaluate dose-response |
| Coherence | *In vitro* to *in vivo* extrapolation | Experimental evidence from *in vitro* and *in vivo* * |
| Analogy | Predictive (of the outcome of interest) | Selection based on the KEs preceding the AO of interest |
| Specificity | Specificity | Gene ranking based on the specificity score |
| Plausibility | (Biological) plausibility | The AOP framework provides a plausible context; supporting evidence; selection of the organism |
| Temporality | Temporality | Transcriptional alteration follows the exposure; selection of the model organism ** |
| – | GLP-method | RT-qPCR |
| – | Influence | Centrality measures from human protein-protein interaction and gene regulatory networks |

\*The biomarkers selected here are targeted for the development of non-animal assays for toxicological assessment. Hence the coherence to *in vivo* set ups is not evaluated experimentally. However, *in vivo* data was used for the selection of the markers to provide context of the systemic response. \*\*Temporality in the Bradford Hill criteria refers to a clear distinction of the exposure happening prior to the outcome. Here, we considered temporality by observing transcriptional changes post exposure as well as in the selection of the model organism. Macrophages have a crucial role in the initiation of the profibrotic response preceding the outcome, fibrosis.

The prioritisation and selection of the candidate biomarkers considered three main aspects:

1. the social life of genes, i.e., some genes (gene products) are more influential than others,
2. specificity regarding the endpoint of interest, and 3) experimental evidence suggesting the genes respond to a relevant exposure. The ranking of the genes was based on the first two considerations, while the experimental evidence was included to guide the selection of candidate genes for RT-qPCR validation from the ranked list. This enabled a flexible selection process that would be applicable even in the absence of experimental data. At this stage, we also considered the biological feasibility of the target genes given the selected macrophage model as well as a broad coverage over the PF KEs.

As a result, we obtained a list of 25 candidates out of the original 2,075 genes related to PF (Table 2). Although we focused on the genes in the top 10%, we further included genes ranking lower to obtain a broader coverage over the PF KEs. Genes that are specific to individual KEs might rank low when the individual lists are combined. Hence, we considered the expression patterns from the experimental data as well as the specificity scores and ranks in the individual KEs. This also allowed us to evaluate whether genes ranked higher would perform better than others.

**Table 2:**
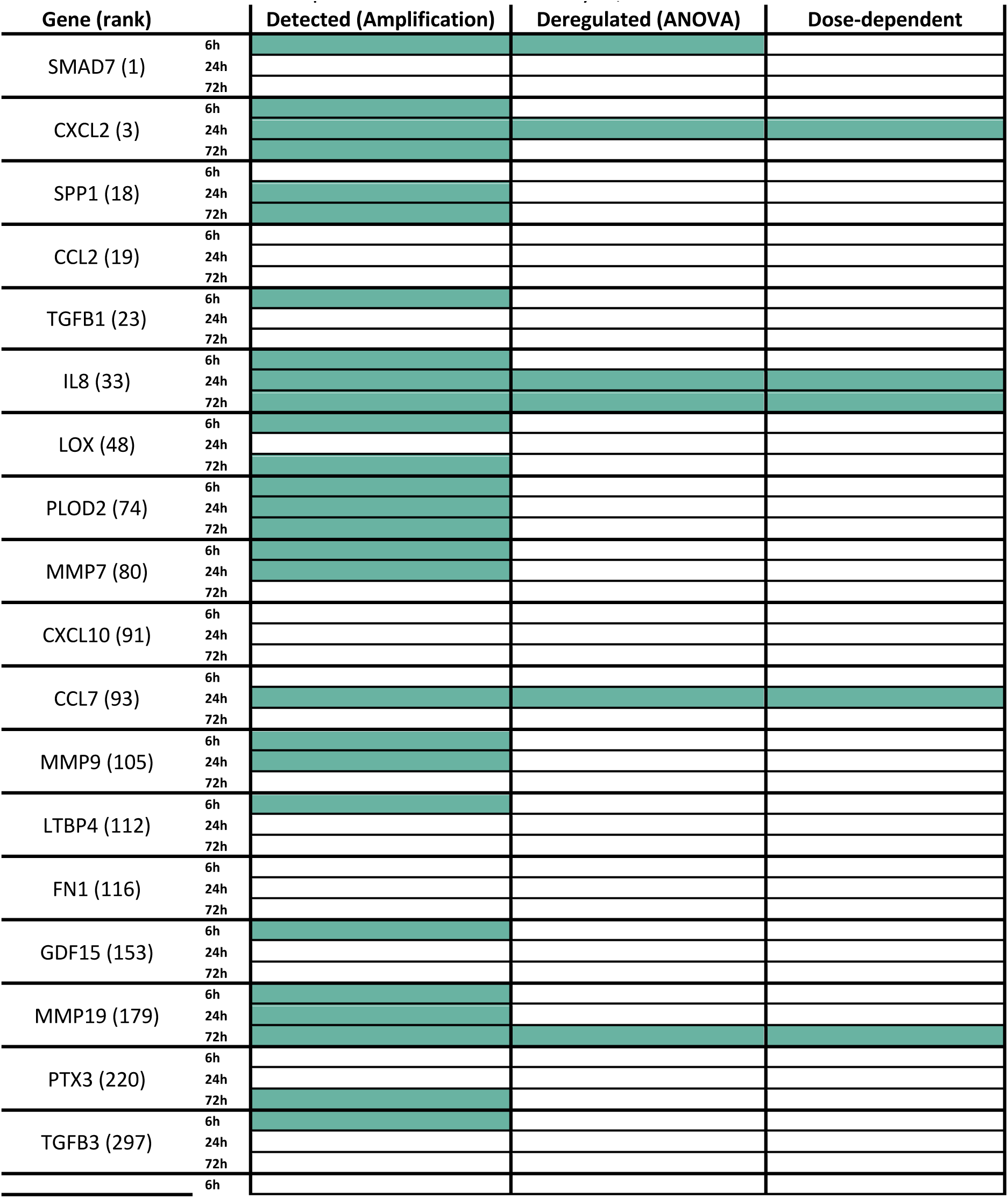
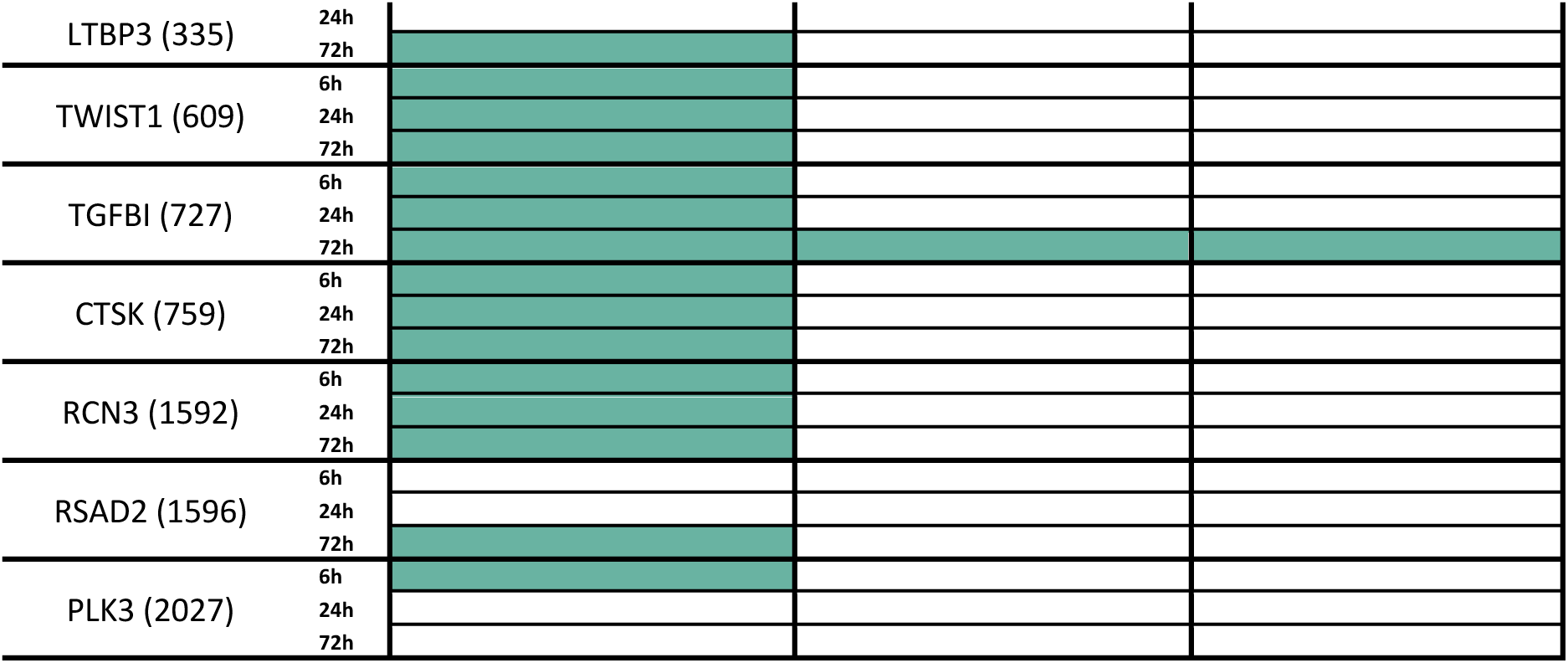
Genes selected for qPCR validation. Green = yes, white = no.

We could detect the expression of 22 of the candidate genes at one or more of the evaluated time points, and six of the detected genes showed significant alteration as compared to the unexposed control samples (Table 2). Finally, five of these genes were altered in a dose-dependent manner: CXCL2 and CCL7 at 24h, IL8 (CXCL8) at 24h and 72h, and MMP19 and TGFBI at 72h. All but TGFBI of these genes were among the top 10% in the global PF rank (Table 2). Although we were not able to fit a dose-dependent model on the highest ranked gene, SMAD7, a suggestive trend could be appreciated in its expression pattern (Figure S4 panel 18/6H). The expression of each of these genes was upregulated as compared to the controls (Figure S4).

The central role of TGF-beta signalling is well-established in PF (76), but neither of the TGFB genes tested (TGFB1 and TGFB3) showed significant change in expression in our setup. Indeed, TGF-beta is activated through a complex cascade of events, where the inactive form of the protein is activated by other effectors post-translationally (77), making members of the TGF-beta family a more attractive target for protein-based biomarker assessment over gene expression. At the same time, we did observe upregulation of SMAD7 and TGFBI which are both activated by TGF-beta (78,79), suggesting the induction of TGF-beta signalling. The protein encoded by TGFBI is involved in the extracellular matrix (ECM), and it has been shown to bind type I collagen, resulting in thicker fibres and further affecting macrophage polarisation towards the M2 type (80). Indeed, bleomycin has been suggested to polarise macrophages towards M2 (often referred to as the anti-inflammatory type), which have been shown to drive the development of PF through their ability to promote myofibroblast differentiation (81,82). Many of our suggested biomarkers are chemokines that mediate immune responses. IL8 and CXCL2 are best characterised as neutrophil attractants, while CCL7 targets a wide variety of leukocytes (83-85). Indeed, prolonged inflammation, combined with persistent M2 macrophage activation, supports pathogenesis of fibrosis (86), and a mixed status of M1/M2 macrophage activation has been previously associated with carbon nanotube -induced PF *in vivo* (87). Similarly, multi-walled carbon nanotubes have been shown to induce the polarisation of macrophages towards such mixed status of M1/M2 polarisation (88,89). MMP19 is a member of the matrix metalloproteinase family involved in ECM remodelling (90). MMPs have been extensively characterised in the context of PF (91,92), and MMP19 specifically has been suggested as a key regulator of PF in mice and humans (93).

Although macrophages alone cannot capture all the KEs of PF, our model is able to highlight the key steps of macrophage involvement in PF. The temporality of the expression of our suggested biomarkers is supportive of the events leading to the development of fibrosis, where the initial inflammation is followed by type M2 macrophage activation that together contribute to the development of a profibrotic microenvironment and responses in other cells in the tissue (86).

New approach methods (NAMs) are urgently needed to reduce animal testing while providing robust evidence to support chemical safety assessment. While alternative methods have been successfully developed to capture acute effects, modelling long-term outcomes of the exposures, such as fibrosis, *in vitro* is still a challenge. Here, we propose a panel of five genes CXCL2, CCL7, IL8, MMP19, and TGFBI as AOP-derived robust biomarkers of PF to be successfully measured in a model of human macrophages *in vitro* after short exposure time.

## Conclusions

Mechanistic toxicology encompasses the means for faster, cheaper, and more ethical chemical safety assessment. However, to unleash the full potential of mechanistic evidence also in the regulatory framework, robust approaches to build confidence towards toxicogenomics are urgently needed. Here, we introduced an integrated approach that links toxicogenomics with the concept of AOPs and proved its applicability to chemical grouping and development of data-driven NAMs. We introduce the concept of AOP fingerprint of a chemical exposure to evaluate potential systemic outcomes through unbiased interpretation of toxicogenomics data. Our results point to a consistent AOP fingerprint of multi-walled carbon nanotubes extrapolated from both in vitro and in vivo experiments. Finally, we identified and experimentally validated a panel of robust AOP-derived *in vitro* biomarkers for pulmonary fibrosis.

## Materials and Methods

### Definition of a Knowledge Graph -based data structure

We established a Knowledge Graph -based data structure by expanding our previously introduced framework, the Unified Knowledge Space (UKS) (94,95). A detailed description and a full list of integrated data sources are provided in the Supporting Methods. The so formed data structure was utilised throughout the study as described in the following sections.

### Annotation of key events

We applied a multi-step strategy comprising natural language processing (NLP) and manual curation to annotate KEs to established gene sets through pathways, phenotypes, and gene ontologies. The annotation strategy is summarised in Figure S2.

#### Computational prioritisation of KE annotations

To match the descriptions of key events and gene sets we developed an NLP pipeline (Figure S2). The pipeline performs several operations to extract the informative terms from both the descriptions of a key event and a gene set that are scored to assess the degree of matching between the two entities. In detail, first, the raw text was converted to lower case and all punctuation symbols were removed. Second, concepts that span multiple words in the text description were replaced by a single word expressing the same concept to strengthen the matching quality (e.g., the concept “positive regulation” was replaced with the single word “upregulated”). Third, the text was split into tokens which are further processed one by one. Fourth, each token corresponding to a stop word in the English language was dropped. Stop words refer to the most common words in a language that do not bring additional meaning (e.g., for the English language common stop words include “in”, “the”, “of”, “from”). Fifth, different declinations of the same concept were mapped to their root term (e.g., plurals were converted to singulars, the terms “increased” and “increasing” were both mapped to “increase”). We also used this same procedure to standardise several styles to write the same symbol (e.g., “pparα” and “pparalpha” map both to “ppar-alpha”). After these preprocessing steps, each gene set and key event was represented by a set of token words, e.g. {upregulate, ppar-alpha}. However, the frequency of each token word across the descriptions of genes and key events is not the same, and hence, the informative value of rare terms is higher than the informative value of more common tokens. We took this into account by weighting each token by its inverse document frequency (IDF), i.e., the weight was inversely proportional to the number of gene sets and key events that contain that token. Finally, we employed a weighted version of the Jaccard Index (JIW) to match gene sets and the key events, using the IDF as weights (i.e., each token that was shared between a gene and a key event did not account 1 as in the standard Jaccard Index, but it contributed its IDF weight to the matching score) and sorted the matching gene sets for each key event in descending order.

#### Manual curation and refinement of annotations

Next, the results of the computational prioritisation were manually evaluated for correct context and accuracy. Manual curation was used for gap filling and refinement of the annotations. In detail, the top five matches retained from the natural language processing -based approach were evaluated, and inaccurate or spurious matches were discarded. In case no matches from the computational prioritisation were deemed suitable, a manual search related to the name of the KE was performed on relevant databases (WikiPathways (96), HPO (97), KEGG (98), Reactome (99) and GO (100)). For molecular level KEs, where the alteration of an individual gene was described, either the main functions of the gene were selected, or the gene was directly annotated to the ensemble identifier of the said gene. More generic annotations (i.e., annotation of a KE describing the alteration of a gene to a functional term tightly related to that gene) were prioritised to increase the size of the relevant gene sets.

The matches for KEs expressing the increase/activation or decrease/repression of a biological progress were further organised based on the hierarchy of the terms by prioritising the most generic but suitable term followed by increasing specificity when multiple annotations of various specificities were available. For instance, Ke:1457 called “Induction, Epithelial Mesenchymal Transition” was annotated to the following terms: 1) Epithelial to mesenchymal transition (GO:0001837), 2) Regulation of epithelial to mesenchymal transition (GO:0010717) and 3) Positive regulation of epithelial to mesenchymal transition (GO:0010718). The curated KE – gene set links were added to the UKS so that for each key event entity its top five matches were added, while the matching level was stored as an edge attribute. This allows to either combine multiple mappings for a key event or to filter for specific mapping levels. Since the KE – gene set mappings are always the same for the same KE, these relationships were added to the *Key Event* entities and not to the *Specific Key Event* entities, which reduces complexity of the knowledge graph as well as reduces needed storage space. The information, however, can still be retrieved from the UKS via its connecting paths.

#### Gene set retrieval

The genes corresponding to the matched terms were retrieved by matching the term names to their exact identifiers and querying the UKS for human genes associated with the terms. For phenotypes (HPO and KEGG disease), only genes with a link in the original database were included by filtering by the source for the connection. In cases where no human genes were linked to the annotated GO term, we obtained the mouse and rat genes associated and converted them to human orthologs using Ensembl (101), which were then used as the corresponding gene sets. When no genes of human, mouse, or rat were associated with the original term, the annotation match was discarded and considered unsuccessful. Once gene sets to all original terms were defined, the gene sets were merged to obtain the final set of genes corresponding to each KE in this study.

### Enrichment analysis of reference chemical -associated gene sets

To evaluate the ability of our framework to highlight relevant adverse outcomes from chemical associated gene signatures, we retrieved lists of reference chemicals from the EU Reference Laboratory for alternatives to animal testing (ECVAM) reference chemical library (102) and National Toxicology Program Interagency Center for the Evaluation of Alternative Toxicological Methods (NICEATM) website (https://ntp.niehs.nih.gov/whatwestudy/niceatm/resources-for-test-method-developers/refchem/index.html). From the resources provided by ECVAM, we selected a hepatotoxic chemical list that had clear distinctions between positive and negative compounds. This list was based on the work from EPA’s Virtual Liver project (https://cfpub.epa.gov/si/si_public_record_report.cfm?dirEntryId=166616&Lab=NCCT), and was provided as a downloadable Excel-file (./CHELIST/CheLIST EPA_VLIVER.xlsx) by ECVAM. From NICEATM, we selected the list of chemicals with characterised thyroid activity (specified as “ACTIVE” in the listing produced based on a previous publication by Wegner et al. (103). Androgen receptor and estrogen receptor agonists were selected from the lists of *in vitro* reference chemicals provided on the website. These listings had been previously published in Kleinstreuer *et al*. (104) and Browne *et al*. (105), respectively. Finally, carcinogenic compounds were identified from the list containing chemicals that are either known carcinogens or reasonably anticipated to be human carcinogens (RAHC) based on the 14^th^ report on Carcinogens (RoC Classifications) provided by NICEATM. The chemicals from each of the reference lists were then matched to the list of chemicals obtained from the Comparative Toxicogenomics Database (CTD) (29) though name-based matching or by the provided CAS identifiers, resulting in the final lists of reference chemicals for each endpoint used in this study. Chemical-gene links originating from the CTD were retrieved from the UKS and only chemicals with 50-1,000 associated genes were included in the enrichment analysis.

Enrichment analysis was performed using the Fisher’s exact test as implemented in the function *enrich* from R package bc3net (106) for each chemical associated gene set against the list of AOP-related genes (*i.e.*, the union of all the genes associated to all the KEs of the AOP). Enrichment p-values were adjusted using the false discovery rate (FDR) correction. AOP was considered significantly enriched with FDR-corrected p-value < 0.01.

### KE clustering and construction of the pulmonary fibrosis network

Similarities between the gene sets associated to each KE were evaluated by calculating the Jaccard Index (JI) between all pairs of KEs (size of the intersection divided by the size of the union of the gene sets). The resulting similarity matrix was then used to group the KEs using hierarchical clustering as implemented in the function *hclust* in R package stats, specifying the agglomeration method as “ward”. The number of clusters was defined so that only KEs with the same gene sets associated to them (JI = 1) were assigned to the same group. The grouping obtained in this manner was used to perform the enrichment against KEs to avoid multiple testing against the same gene set as well as to enhance the network presentation of the pulmonary fibrosis (PF) AOP network. The unweighted PF AOP network was generated using gephi (107) by importing a graphml file generated with the function *graph_from_edgelist* from R package igraph (108). KE groups from the clustering were added as attributes to the nodes and used to merge redundant nodes in gephi. Similarly, AOPs each KE is associated to were added as attributes and used to colour the nodes.

### Characterisation of the AOP-fingerprints

#### Transcriptomics data

*In vivo* and *in vitro* transcriptomics data from MWCNT (Mitsui-7) exposures were selected from a previously published collection by Saarimäki et al. (67). The original data sets are available under GEO accession number GSE29042 (*in vivo*) and ArrayExpress entry EMTAB6396 (*in vitro*), while the preprocessed data is available on Zenodo (https://doi.org/10.5281/zenodo.6425445). The *in vivo* data set comprises multiple doses and time points, while the *in vitro* data contains a single dose and time point exposure on four distinct cell lines representative different cell types of the lung. In each case, differentially expressed genes (filtered by an absolute fold change > 1.5 and Benjamini & Hochberg adjusted p-value < 0.05) obtained from Zenodo (https://doi.org/10.5281/zenodo.6425445) for each distinct comparison (i.e., combination of each dose and time point vs. control *in vivo* and separate cell lines *in vitro*) were pooled together to obtain a distinct mechanism of action of the exposure *in vivo* and *in vitro*, respectively.

#### AOP fingerprints

To produce the AOP fingerprint for the MWCNT exposures, enrichment analysis was performed using the Fisher’s exact test as implemented in the function *enrich* from R package bc3net (106) separately against the AOP-associated gene lists and the KE-associated gene lists (KEs linked to the same set of genes were grouped to avoid multiple tests against the same set). An AOP was considered significantly enriched when the AOP itself and at least 33% (or minimum of 2 KEs when the length of the AOP was less than 6) of its KEs were enriched with an FDR-corrected p-value < 0.05.

### Selection and testing of AOP-driven biomarkers

#### Gene prioritisation

All human PPI edges were extracted from the UKS and used to create a robust gene – gene network. PPI/gene (product) interaction information can vary across data sources as well as the covered genes may differ. In addition, there is an innate bias in the data, where more data sources are available for “more investigated” genes and gene products. Because of this, we decided not to apply a global threshold on how many sources need to support an edge (109), but instead, we applied a local threshold. This ensures that also less investigated genes and gene products will be retained in the final robust gene – gene network, but their edges are less penalised by the number of supporting edges, than highly covered gene (product) nodes. For each node, the mean number of sources supporting all its connecting edges was estimated and only edges with at least the “mean number of sources” for a node were retained, which needed to be true for at least one of the nodes making up an edge. This was only done for GENE nodes, which were flagged as protein coding in Ensembl. The final robust human gene – gene network, consisted of 20 260 nodes, 806 250 edges, a network density of 0.0039. Due to the significant lower number of available sources for transcription factor – gene (product) data, all available sources were kept and scored equally. These edges were used to create a directed gene – gene network, consisting of 18 754 nodes, 363 649 edges and a network density of 0.001. On the so created gene – gene networks, for each node its degree, betweenness, eigenvector, and closeness centrality were estimated with NetworkX (110). These measures were then used to rank the genes linked to the KEs in the context of individual KEs. The gene list for each KE was ranked based on each of the centrality measures (degree, betweenness, closeness and eigenvector centrality) individually from most central to the least. The ranked lists were combined using the borda method as implemented in the function *Borda* in R package TopKLists (111).

#### Biomarker selection

The gene centrality-based ranking was then supplemented by a specificity ranking for the KEs of AOPs related to pulmonary fibrosis (PF). We calculated a specificity score for the genes in the context of the KEs by dividing the occurrence of the gene in the KEs of PF AOPs by their occurrence in the KEs of other AOPs. Similar score was calculated at the level of AOPs (occurrence in PF AOPs/occurrence other AOPs), as we were looking for universal PF biomarkers (i.e., prioritising those that would be present in as many of the six PF AOPs as possible) while also being as specific as possible to PF. These ranks were again combined by the function *Borda* from R package TopKLists (111), and a final round of the borda method was applied to combine the lists of genes from each KE into one PF rank. We complemented the final rank with experimental evidence. We assessed whether the genes were differentially expressed in the Mitsui-7 exposures *in vivo* and *in vitro*. We also evaluated whether they were dose-dependently altered in the *in vivo* data as well as in an additional *in vitro* data set on Mitsui-7 exposure of PMA-differentiated macrophages (originally published in Saarimäki *et al*. (58) and the preprocessed data available as GSE146708 in the previously published collection (67) available in https://doi.org/10.5281/zenodo.6425445). The dose-response modelling of the *in vivo* (GSE29042) and *in vitro* (GSE146708) datasets was performed by following the strategy implemented in the BMDx tool (112). Particularly, for each gene present in the dataset, multiple models were fitted including linear, second order polynomial, hill, power, and exponential model. For each gene, the optimal model was selected as the one with the lowest Akaike Information Criterion (AIC). Genes with an optimal model with lack-of-fit p-values lower than 0.1 were removed from the analysis. The effective doses (BMD, BMDL, and BMDU) were estimated under the assumption of constant variance and by using a BMRF factor of 1.349 (corresponding to a minimum of 10% of difference with respect to the controls). Genes were further filtered based on the predicted doses. Genes with BMD or BMDU values extrapolated higher than the highest exposure dose were filtered. Moreover, genes whose ratio between the predicted doses is higher than the suggested values (BMD/BMDL> 20, BMDU/BMD> 20, and BMDU/BMDL> 40) were removed from the analysis. Genes passing the filters were considered to be dose-dependently altered. At this stage, we also considered the measurability and feasibility of the gene in the selected macrophage model. For instance, numerous collagen-encoding genes were ranked high, but would not be a meaningful target in a macrophage model. Moreover, we aimed for a high coverage of PF KEs and aimed to select genes with high specificity scores. With these considerations, we selected a subset of the genes with the following priority: 1) genes that are deregulated both *in vivo* and *in vitro*, with most emphasis on dose-dependency, 2) genes that are deregulated *in vitro*, with most emphasis on dose-dependency and 3) genes that are not significantly differentially expressed but are dose-dependent. Finally, after this initial selection driven by the rank and experimental evidence, we included additional candidate biomarkers that had a lower rank but were specific to KEs that would otherwise not have been covered by the selected candidates.

#### Cell culture

THP-1 cells (DSMZ no.: ACC 16) were grown in RPMI 1640 (Gibco, #21875) + 10% inactivated FBS (Gibco, #10270). Cells were cultivated in 75cm^2^ culture flasks at 37°C with a humidified atmosphere of 5% CO2. For all experiments, cells were seeded at a density of 1x10^5^ cells/ml in 96 well plates and differentiated for 48 hours with 25 nm PMA (phorbol-12-myristate-13-acetate, Sigma-Aldrich, #P1585). Cells were then left to rest for 24 hours in fresh media containing no PMA prior to bleomycin exposures.

#### Cell Viability Assay

THP-1 cells were exposed to 0-10 µg/ml of bleomycin ready-made solution (Sigma-Aldrich, #B7216) and 0-100 mg/ml of Triclosan (Sigma-Aldrich, #72779), for 6, 24 and 72 hours. A WST-1 assay was then used to measure cell viability. Briefly, 10 µl of cell proliferation reagent WST-1 (Roche, #11644807001) was added to each well. Cells were left to incubate with WST-1 for 3 hours in a 37°C, 5% CO2 incubator. Absorbance at 450 nm was then measured with a Spark microplate reader (Tecan). Results of the cell viability assay are available in Supplementary file 2 and Figure S3.

#### RT-qPCR

For each time point of 6, 24 and 72 hours, THP-1 cells were exposed to 0, 2.5, 5, 10 and 100µg/ml of bleomycin ready-made solution (Sigma-Aldrich, #B7216). Media was removed and cells were washed briefly with 50 µl of PBS. 100 µl of lysis buffer from the QIAGEN RNeasy mini kit (Qiagen, #74104) was added to each well to lyse the cells. 3 wells (300 µl) were pooled to create one sample, and there were 5 samples for each concentration at each time point. Total RNA was then extracted from these samples using the QIAGEN RNeasy mini kit (Qiagen, #74104). cDNA was synthesised from 100 ng of RNA, using the High-capacity cDNA Reverse Transcription Kit (Thermo Fisher Scientific, #4368813), according to manufacturer’s instructions. Expression levels of target genes were determined by qRT-PCR using CFX96 Touch Real-Time PCR Detection System (BioRad) with 10 µl of iQ Multiplex Powermix (Bio-Rad, #1725849), 5 μl of cDNA diluted 5-fold, 2.5 µl of Nuclease-free (NF) water (not DEPC-Treated, ThermoFisher, #AM9930) in a 20 μl reaction, together with 2.5 µl of single (1 µl assay + 1.5 µl NF water) or multiplexed (0.5 µl of each assay) PrimePCR Probe Assays (Bio-Rad) as follows with single or multiplex reactions grouped in parentheses and formatted as *Gene*/UniqueAssayID: (*ACTB*/qHsaCEP0036280), (*SMAD7*/qHsaCEP0050142, *MMP9*/qHsaCIP0028098, *GDF15*/qHsaCEP0051579, *CTSK*/qHsaCIP0030907, *PLOD2*/qHsaCEP0052848), (*CXCL2*/qHsaCEP0058163, *LTBP4*/qHsaCEP0024931, *TGFB3*/qHsaCEP0058244, *RCN3*/qHsaCEP0057804, *MMP7*/qHsaCEP0052037), (*SPP1*/qHsaCEP0058179, *FN1*/qHsaCEP0050873, *LTBP3*/qHsaCEP0053782, *RSAD2*/qHsaCIP0031596, *CCL7*/qHsaCEP0058033), (*IL8*/qHsaCEP0053894, *MMP19*/qHsaCEP0051244, *TWIST1*/qHsaCEP0051221, *PLK3*/qHsaCIP0027687, *CXCL10*/qHsaCEP0053880), (*LOX*/qHsaCEP0050731, *PTX3*/qHsaCEP0033071, *TGFBI*/qHsaCEP0058394, *CCL2*/qHsaCIP0028103, *TGFB1*/qHsaCIP0030973).

Fold change (FC) values from RT-qPCR data were calculated using the comparative CT(2^−(ddCt)^) method (113). The FC values were log2 transformed (log2(FC)). For each gene and for each concentration, an outlier detection was performed by removing all the samples with log2(FC) values above or below the 75^th^ and 25^th^ percentiles of the distribution. Ct values, dCt values, FC values and log2(FC) values are available in Supplementary file 2 along with ANOVA tables and tukey HSD posthoc test results. The log2FC expression of the genes as compared to the untreated controls are plotted in Figure S4.

#### Dose-dependent modelling

A dose-response analysis of the log2(FC) values derived from the PCR experiments was performed. For each gene, multiple models were fitted, including linear, hill, power, polynomial, exponential, log-logistic, Weibull, and Michaelis-Mentel models. The optimal model was selected as the one with the lowest AIC. The BMD estimation was performed under the assumption of constant variance. The BMR was identified by means of the standard deviation approach with a BMRF of 1.349. Only genes with lack-of-fit p-value >0.10 and with estimated BMD, BMDL and BMDU values were considered relevant.

## Data Availability

The full list of KE annotation is available from the corresponding author upon request.

## Acknowledgements

This work received funding from the EU Horizon 2020 project NanoSolveIT (grant agreement no. 814572) as well as the Academy of Finland (grant agreement no. 322761). Saarimäki was supported by the Emil Aaltonen Foundation.

